# Distinct resource utilization by introduced man-made grouper hybrid: an overlooked anthropogenic impact from a longstanding religious practise

**DOI:** 10.1101/2023.10.12.562133

**Authors:** Arthur Chung, Celia Schunter

## Abstract

Anthropogenic activities, such as non-native aquaculture species introduction have been considerably altering trophic interactions in marine ecosystems. The hybrid grouper (TGGG), an aquaculture product originated from artificial F1 crossbreed between *Epinephelus fuscoguttatus* and *E. lanceolatus*, have been released in the wild through religious activities, raising concerns about this man-made introduced species. The carnivorous diet, together with large body size inherited from the parental species have made TGGG a candidate that could pose significant impacts to the marine ecosystem. Yet, little is known about the diet composition of TGGG upon release into the natural environment, and any dietary overlap or partition with closely related endemic species. Here, we deploy gut content DNA metabarcoding to determine the prey richness and dietary niche of wild caught TGGG and compare with four native grouper species (*Epinephelus awoara, E. bleekeri, E. coioides and E. quoyanus*). The TGGG exhibited six unique prey taxa with teleosts being the major taxa preyed upon followed by crustaceans and cephalopods and displayed significant lower mean number of pray taxa compared to other groupers. The TGGG exhibits a significantly different diet composition, possibly indicating a diet transitioning and acquiring new feeding behavior. This study provides a comprehensive analysis with high taxonomic resolution on the diet of artificial hybrids in the wild, suggesting possibility of introduction success if release events persist. Finally, these findings provide new information on how local trophic dynamics are impacted by under-investigated release of animals through religious practices.

## Introduction

The study of trophic interactions is fundamental to understand species’ roles in the food web and the interactions that shape the complex dynamics in trophic networks (Kéfi et al., 2012; Parravicini et al., 2020). In particular, prey assembly structure in consumer stomach content can disclose ecological mechanisms, including intra-specific competition for resources, expansion of dietary niche at individual to population level, and inter-specific niche partitioning through divergence in morphology and behavior (Colloca et al., 2010; Ratcliffe et al., 2018; Varghese et al., 2014). With anthropogenic activities considerably altering trophic interactions in marine ecosystems (Gilarranz et al., 2016; Johannesen et al., 2012), dietary studies provide empirical evidence on changed predator-prey relationships, for instance, due to climate change (Davoren et al., 2012; McMahon et al., 2019), ocean acidification (Poore et al., 2013; Watson et al., 2017) and overfishing (Audzijonyte et al., 2013; Roux et al., 2013).

Coastal aquacultural practices also have significant effects on marine ecosystem dynamics (López et al., 2008; Ferriss et al., 2016). Notably, farm escapes where release of farmed species leads to competition for resources and predation on local fauna in the wild (Toledo-Guedes et al., 2014; Valero-Rodriguez et al., 2015). Such farm escapes are also responsible for other adverse ecological impacts such as transmission of pathogens and pollution of the local gene pool through interbreeding or hybridization (Arechavala-Lopez et al., 2018). An additional yet understudied way of introduction of aquaculture species into the wild is by mercy release. In this Buddhist or Taoist religious practice, also known by alternative names such as prayer animal release and religious release, captive animals are released to gain spiritual merit (Wasserman et al., 2019). Mercy release in marine or coastal environment is prevailing in Asia, including Hong Kong, Taiwan and Singapore (Crook, 2023; Jaafar et al., 2012; Shea & To, 2018). Such practices commonly involve release of numerous non-native species with high potential of biological invasion (Everard et al., 2019; Magellan, 2019).

The hybrid grouper (TGGG), a F1 hybrid species developed artificially from two large parental grouper species *Epinephelus fuscoguttatus* and *E. lanceolatus* (Ch’Ng & Senoo, 2008), is both a highly demanded aquaculture product and a popular choice for mercy release (Cheang et al., 2020). TGGG can grow to enormous size with reported total length longer than 85cm and weight up to 20kg in a growth rate surpassing both parents (Ching et al., 2018; Luin et al., 2014; Shapawi et al., 2019). TGGG is also capable of developing mature gonad and reproduce through natural spawning that give rise to viable F2 offspring (Ching et al., 2018; Luin et al., 2014). These factors, combined with uncontrolled mercy release practices has made TGGG a potential candidate that could pose significant impacts to the marine ecosystem.

However, effects of TGGG on the receiving ecosystem have never been evaluated. Examples from farm escapes of aquaculture species, such as the rainbow trout *Oncorhynchus mykiss* suggests that hatchery individuals are capable of consuming a large variety of prey in the wild, some even indistinguishable from wild occurring rainbow trout, with diet also depends on season, size and dispersal distance from the originated fish farm that reflect a diet transition from aquaculture to wild (Nabaes Jodar et al., 2020; Rikardsen & Sandring, 2006). *O. mykiss* farm escapes was also found to have a broader dietary breadth than wild rainbow trout, although the diet included indigestible items as well (Nabaes Jodar et al., 2017). Yet, the diet of TGGG cannot be measured and compared in the same way as *O. mykiss* farm escapes, as there is no wild TGGG hybrids to allow direct dietary comparison and interpretation of differences of diet. Another way to assess prey preference of a hybrid would be to analyse the diet composition of the parental species which for a coral reef fish hybrid revealed a high diet overlap with both parental species in the same reefs (Montanari et al., 2014). However, although both parental species of the TGGG, *E. fuscoguttatus* and *E. lanceolatus,* cover a wide distribution range across the Indo-Pacific, it is rare to find one or both of them co-occurring with TGGG due to a drastic population decrease for both parental species (Sadovy de Mitcheson et al., 2020) .Therefore limited parental dietary information can be provided to infer the diet preference for TGGG. The absence of both parental species from overlapping regions of their natural habitats and mercy release sites also suggested that TGGG could exploit unfilled ecological niche that previously occupied by parental species (Thuiller et al., 2010).

The TGGG thrives in the wild in Hong Kong and has been seen to inhabit similar habitats as native *Epinephelus* species such as coral communities and nearshore boulder substrates (Shea & To, 2018). It is therefore speculated that TGGG could share similar trophic resources and compete with its closely related grouper species. *Epinephelus* spp. are generally considered as non-specialized carnivores with diets that include fish, crustaceans and zoobenthic animals. Yet, despite the abundance and diversity of species within the genus in the wild, the dietary information and food resource partitioning between the sympatric *Epinephelus* species is not well quantified, other than the feeding ecology studies of one or two species at a time (Chuaykaur et al., 2020; Freitas et al., 2017). Examples of other major carnivorous reef fish species, such as Lutjanidae and Lethrinidae revealed diet partitioning between species utilizing the same reefs (Kulbicki et al., 2005; Takahashi et al., 2020), suggesting that *Epinephelus* species could potentially display a similar dietary resource division. With a deeper understanding of dietary diversity and trophic niche partitioning of *Epinephelus* species, we can discover the range of ecological roles and dynamics of TGGG and compared other grouper species within the same ecosystem.

In this study, we utilize a multi-gene metabarcoding assay approach to perform dietary analyses on TGGG and four native groupers, namely *Epinephelus awoara, E. bleekeri, E. coioides and E. quoyanus* (Figure 1). These species are abundant in Hong Kong waters and share similar habitats along the coast (Sadovy & Cornish, 2000), making them ideal candidates to characterize the diversity of prey and allow for a baseline for dietary comparison and potential partitioning with the TGGG hybrid. Our multiple (gene) taxon-specific assay approach provides a comprehensive coverage of ingested prey while reducing issues of primer specificity and bias (da Silva et al., 2019). We hypothesize that 1) dietary partitioning exist among the four native *Epinephelus* species as indicated from other families of carnivorous reef species, and 2) TGGG exhibit a broader, yet overlapping dietary niche when compared with native species due to diet transitioning and variation between released individuals such as characterized by other farm escapes. By evaluating the diets of these key predators through metabarcoding we enhance our understanding towards ecological resources utilization and evaluate the potential ecological impacts from introduction of an artificial hybrid into marine communities.

**Figure 1.**
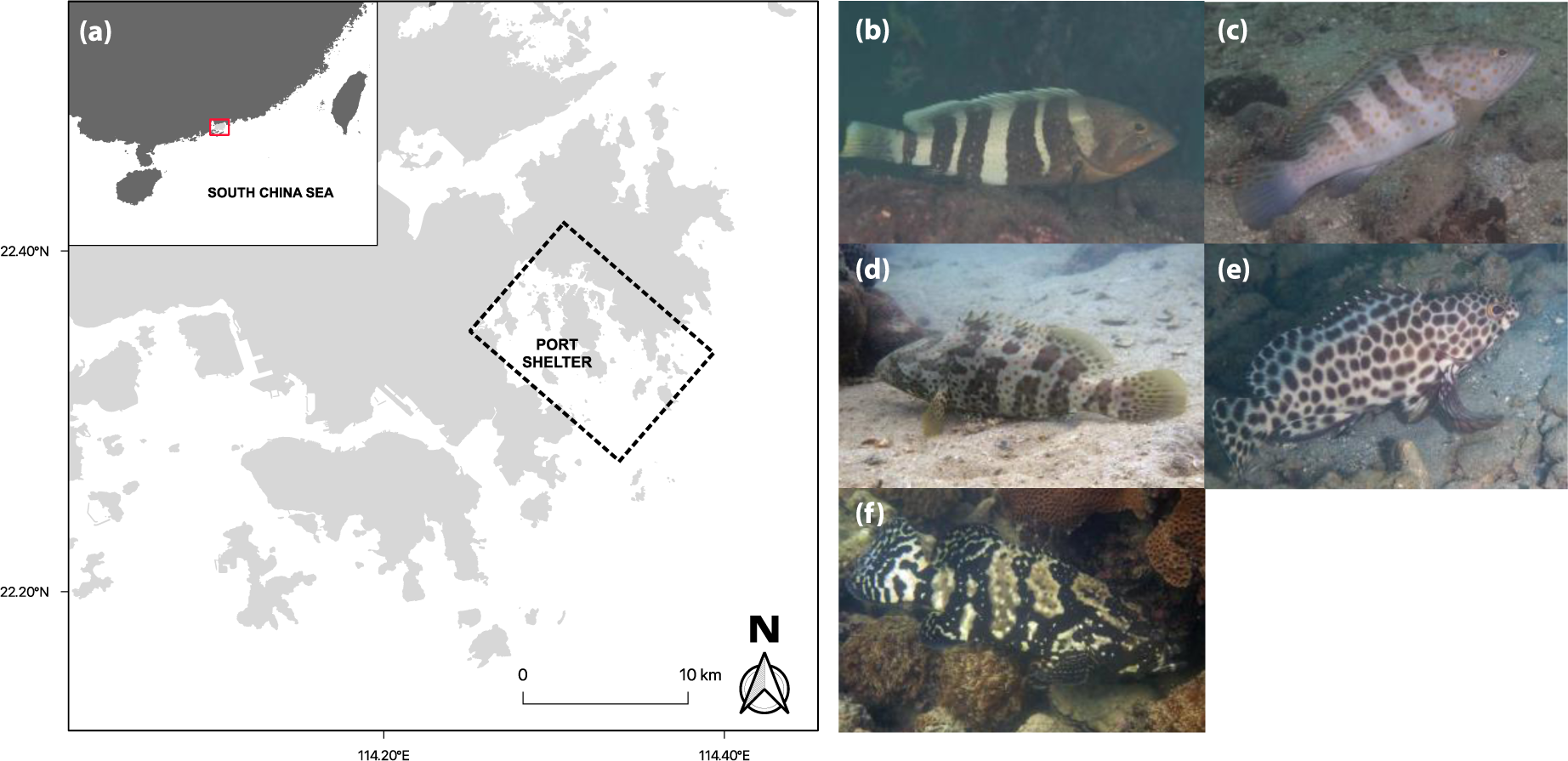
(a) Map of the study area in Hong Kong. The sampling site, Port Shelter Bay is highlighted in black dashed box. The inset on upper left shows location of Hong Kong within the South China Sea. Photos on the right are the species collected in this study, namely (b) *Epinephelus awoara*, (c) *E. bleekeri*, (d) *E. coioides*, (e) *E. quoyanus* and lastly (f) *E. fuscoguttatus* x *E. lanceolatus* (TGGG). Photographs: Allen To, Eric Keung, Ryan Cheng, Stan Shea and YiuWai Hong from 114°E Hong Kong Reef Fish Survey.

## Methods

### Sample collection and processing

All four grouper species *Epinephelus awoara, E. bleekeri, E. coioides, E. quoyanus*, and hybrid grouper TGGG (Figure 1b-f) were collected within the confined area of Port Shelter Bay, south of Sai Kung Peninsula in Hong Kong (Figure 1a) to limit potential bias that could be introduced from spatial variation of diet composition among groupers. This area is also found to be one of the most popular locations for mercy release activities, making it a suitable sampling site to investigate hybrid TGGG release. Samples were collected in two collection periods, first during the dry season from October 2020 to February 2021, and second during the wet season from May 2021 to September 2021. *E. awoara* were collected only in the dry season, while *E. quoyanus* were collected only in wet season due to seasonality in fish catches for these species. Several methods were applied for collection to allow for adequate sampling size including recreational spearfishing, recreational angling and commercial fishing with cage trap. Fish catches from recreational methods were first stored in −20°C freezer in individual bags and catches from commercial fishing were first stored on ice and then place into −20°C on the same day of sampling. All samples were then stored whole in the −20°C until dissection.

In sterile laboratory conditions, the fish were thawed, and only intestinal contents were dissected and isolated from fish samples to avoid potential contamination from non-prey DNA sources such as seawater intrusion or bait in the stomach. All dissection tools were cleaned with 70% ethanol, 10% bleach and lastly rinsed with Milli-Q water between each sample to avoid cross-sample contamination. Stomach digestion levels modified from To (2009) that separated into four category were recorded from each sample: 1) little or no digestion of prey except superficially, for example, skin and fins; 2) moderate digestion of prey with head and tail mostly digested and possibility of parts broken off with oval fleshy remains; 3) major digestion of prey with small fleshy remains and abundance of broken parts, and 4) complete digestion of prey with very small fragments or prey remaining or empty stomach and clean lining. All isolated intestinal content was preserved with 95% ethanol and stored in the −80°C freezer. Fin clips were also obtained from each sample for later species identification verification.

### Species and hybrid verification

To ensure the identity of the collected specimen, we collected fin clips from each and extracted DNA using DNeasy Blood & Tissue Kit (Qiagen) following manufacturer’s instructions. Quality and quantity of the DNA were measured with a Nanodrop 1000 spectrophotometer (Thermo Fisher Scientific). PCR was carried out with primers sets that amplified the mitochondrial cytochrome oxidase subunit I (COI) (FishF1: TCAACCAACCACAAAGACATTGGGAC; FishR1: TAGACTTCTGGGTGGCCAAAGAATCA; Ward et al., 2005). TGGG samples underwent another PCR run to amplify the nuclear RYR3 gene region (F48: TGACAGCTTCAGAGAGTATGACCCT; R849: GCCAGTGAAGAGCATCCAGAAGAAG; Qu et al., 2018) in order to identify both parental species of the hybrid (Qu et al., 2018). The PCR mix were prepared as follow: 10μl of Taq PCR Master Mix Kit (Qiagen), 1μM of each primer, 2μl of extracted DNA and bring the total reaction volume to 20μl with PCR grade water. All PCRs were run under following condition: initial denaturation at 95°C for 5 minutes, followed by 35 cycles of 95°C for 30 seconds, 53°C for 30 seconds and 72°C for 45 seconds, with a final extension of 10 minutes at 72°C. The PCR products were visualized by gel electrophoresis on 2% agarose gel and 100bp ladder (SMBIO Technology) and Sanger sequenced at the Beijing Genomics Institute (BGI). The sequencing data were aligned and analyzed using Geneious v11.1.4 and the built in BLAST function. All samples had above 99% identity with the corresponding species.

### Blocking primer development

In order to minimize preferential amplification of the host DNA over prey DNA during polymerase chain reaction (PCR) due to higher concentration of host than prey DNA, blocking primers specific to each target host species were designed following processes described in Takahashi et al. (2020). Briefly, each species-specific blocking primer ordered from Integrated DNA Technologies (IDT; Singapore) was made of 1) 10 bases of 3’ end of the Fish 16S reverse primer; 2) 15 bases following to the reverse primer sequences that were specific to the host species; and lastly 3) a 3-carbon chain (C3) modification at the end of the 3’ end of the blocking primer to inhibit further nucleotide extension (Table 1). Blocking primers were tested with *in-silico* experiments where primer sequences were aligned with 16s sequences of local potential preys that are available through National Center for Biotechnology Information’s (NCBI) GenBank nucleotide database to look at numbers of mismatch bases. The primers were then further tested with two *in-vitro* experiments. Firstly, the optimal blocking primer concentration (100μM and 200μM) and annealing temperature (54°C and 58°C) were tested with qPCR respectively with fixed amount of host genomic DNA (100ng) and the following reaction mix: 1 U of AmpliTaq Gold DNA polymerase (Thermo Fisher Scientific), 1x of AmpliTaq Gold Buffer (Thermo Fisher Scientific), 2mM MgCl2, 0.2mM each dNTP, 0.4 μM of each Fish 16S primer, 0.12x SYBR Green (Thermo Fisher Scientific), 8μg bovine serum albumin (Thermo Fisher Scientific), 100μM or 200μM of respective blocking primer and bring the total reaction volume up to 25μl with PCR grade water. Secondly, the blocking primers were further tested to look for any potential inhibition effects on prey DNA by running a second qPCR experiment with the optimal blocking primer concentrations and annealing temperature verified in the first experiment. Target prey genomic DNA was added in the same reaction mix as above and respective Cq values between treatments with blocking primers added and without were compared. Four species of potential prey from different families were selected for the test, namely *Bathygobius fuscus* (Gobiidae)*, Chaetodon wiebeli* (Chaetodontidae)*, Ostorinchus fasciatus* (Apogonidae)*, Parupeneus indicus* (Mullidae) *and Stethojulis Terina* (Labridae). These selected species and their respective family are commonly found in Hong Kong reef communities. Each qPCR was run on a CFX96 Real-Time PCR Detection system (Bio-Rad) under the following condition: initial denaturation at 95°C for 5 minutes, followed by 50 cycles of 95°C for 30 seconds, 30 seconds of annealing temperature (54°C/58°C) and 72°C for 45 seconds, with a final extension of 10 minutes at 72°C.

**Table 1.**
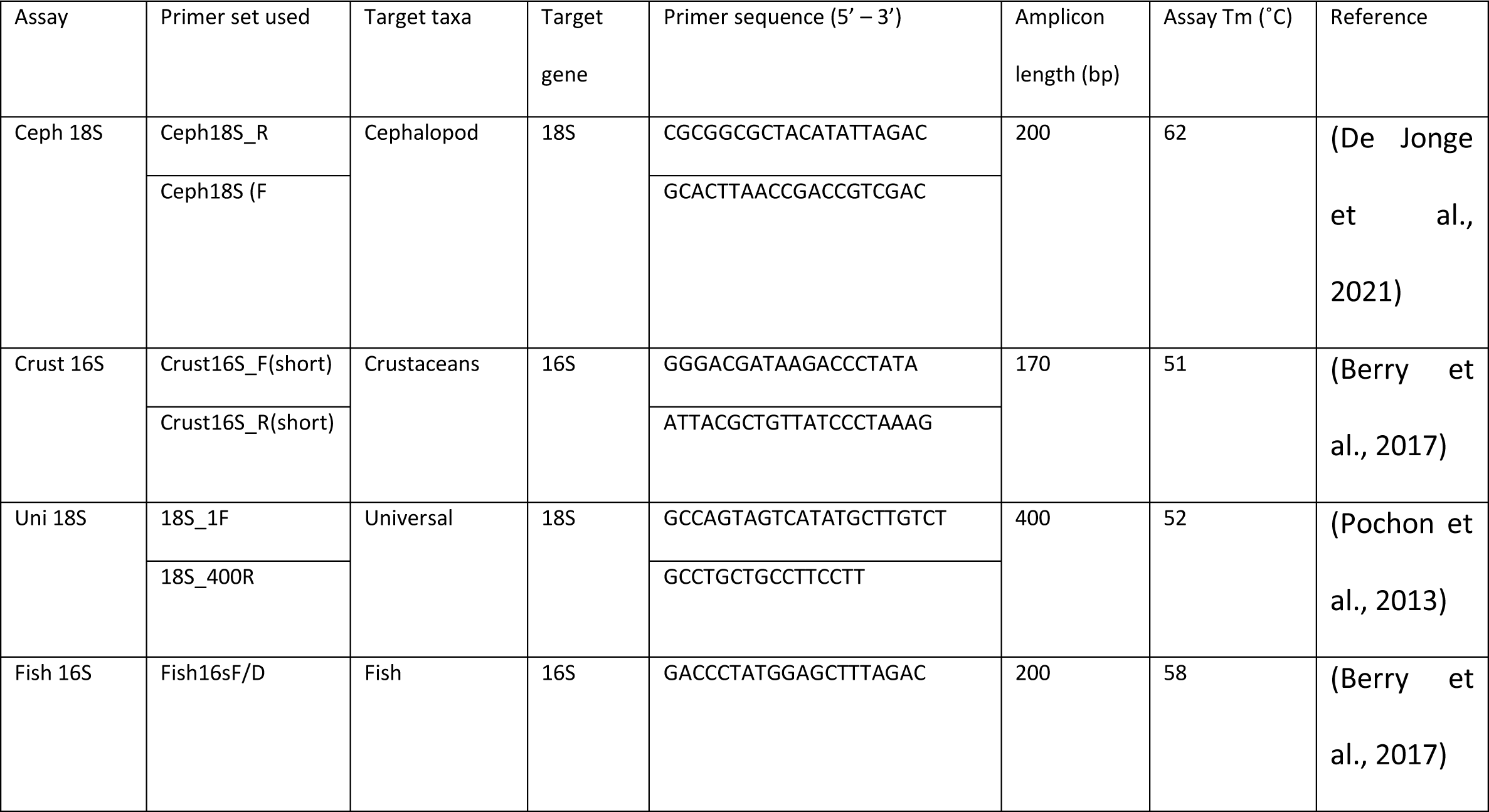
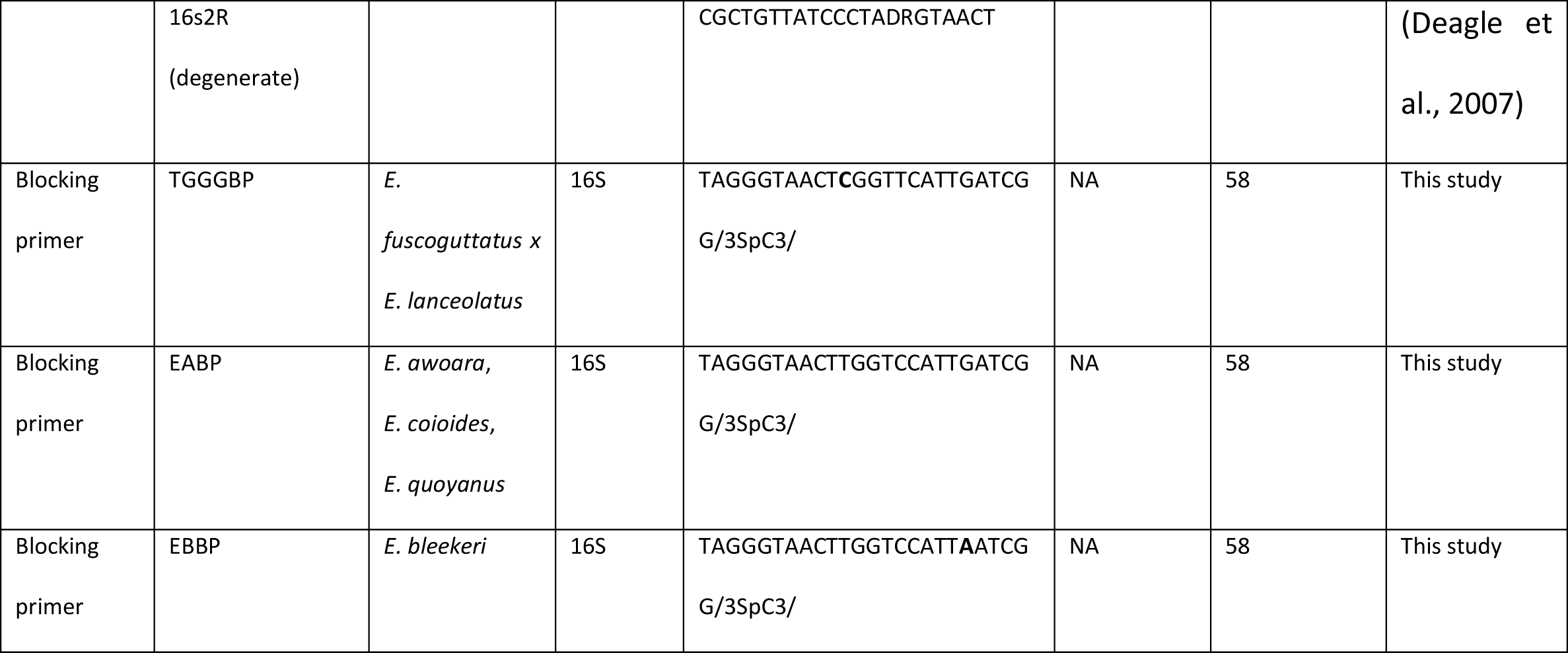
Taxa-specific primer assay used for targeting potential prey items in host species, and associated host-specific blocking primer sequences designed in this study.

### Intestinal content DNA extraction

All steps and procedures of DNA extractions were performed under sterile molecular laboratory conditions. Intestinal contents of fish samples were first homogenized with Tissue Master Lab Homogenizer (Omni) in low speed for 30 seconds. 2ml of the homogenate material of each sample was centrifuged for 5 minutes at 15,000g. The ethanol supernatant was removed and weighed to ensure adequate starting material for extraction. Samples below 60mg received a second round of homogenization and centrifugation. The final weight of samples was between 64mg and 240mg. The pellets were then extracted with QIAamp® PowerFecal Pro DNA Kit (Qiagen) following instructions from manufacturer, with addition of 65°C incubation for 30minutes before the first centrifugation step, and pause the Powerlyzer 24 Homogenizer for 5 minutes instead of 30 seconds between two cycles during the homogenization step. Extracted DNA was measured with a Nanodrop 1000 spectrophotometer.

### DNA amplification and high-throughput sequencing

All steps of PCRs and library preparation were performed inside the safety cabinet with all materials exposed to minimum 15 minutes of UV beforehand. The extracted DNA was first tested for amplification efficiency and inhibition for each assay (Table 1) by qPCR on neat, 1/10 and 1/20 dilutions, which had been shown to favor the generation of accurate species profile from the metabarcoding data (Murray et al., 2015). The amplification mixture contained 1 U of AmpliTaq Gold DNA polymerase (Thermo Fisher Scientific), 1x of AmpliTaq Gold Buffer (Thermo Fisher Scientific), 2mM MgCl2, 0.2mM each dNTP, 0.4 μM of each primer, 0.12x SYBR Green (Thermo Fisher Scientific), 8μg bovine serum albumin (Thermo Fisher Scientific). For Fish16s assay, additional blocking primers specific to each host species with concentration determined in previous experiment were also added to the reaction mix, with total reaction volume adjusted to 25μl with PCR grade water. Each qPCR was run under the following condition: initial denaturation at 95°C for 5 minutes, followed by 50 cycles of 95°C for 30 seconds, 30 seconds of annealing temperature specific to each assay (Represented in Table 1) and 72°C for 45 seconds, with a final extension of 10 minutes at 72°C. The specific DNA dilution with the highest relative template concentration while showing the least inhibited amplification (if any) to a specific assay determined by the Qt values and gel electrophoresis on 2% agarose gel were selected for subsequent metabarcoding using primers with overhang adaptor attached.

The amplicon PCR and attachment of overhang adaptors were performed in duplicate for each sample and respective assay with pre-determined DNA dilution, as well as the PCR mixture and conditions as described above. The size of amplicon products was checked by gel electrophoresis on 2% agarose gel and 100bp ladder. The duplicated amplicons were combined and then purified using the Agencourt™ AMPure™ XP Beads (Beckman Coulter Genomics) following instructions from manufacturer. A dual indexing approach was used to attach sample-specific Illumina sequencing adaptors at both ends of amplicons with the Nextera XT Index Kit (Illumina). The indexing PCR mixture contained 25μl of 2x KAPA HiFi HotStart ReadyMix, 5μl of Nextera XT Index 1 primer, 5μl of Nextera XT Index 2 primer, 5μl of amplicon DNA and bring the total reaction volume up to 50μl with PCR grade water. The PCR was run in the following condition: 95°C for 3 minutes, then 8 cycles of 95°C for 30 seconds, 55°C for 30 seconds and 72°C for 30 seconds and a final 5-minute extension at 72°C. After another round of bead purification as described above, the size and concentration of the indexed products was analysed with D1000 ScreenTape on 4150 Tapestation System (Agilent Technologies), and Qubit 2.0 fluorometer (Thermo Fisher Scientific) respectively. Once the desired peak size and concentration was validated, the libraries were normalized using 10mM Tris-HCI pH 8.5 buffer (bioWORLD) for dilution and pooled together to a concentration of 20nM. The Uni 18S assay were pair-ended sequenced (2 x 300bps) with Illumina Miseq platform (V3 chemistry) due to the amplicon length, while Ceph 18S assay, Crust 16s assay and Fish 16s assay were pair-ended sequenced (2 x 150bps) with illumina Novaseq 6000 platform. All sequencing was performed at the Centre for PanorOmic Siences (CPOS) at the University of Hong Kong.

### Bioinformatic and read filtering

The bioinformatic processes began with demultiplexed sequencing output FASTQ. The primer sequences of each specific assay were then trimmed using Cutadapt v2.8 (Martin, 2011). Corresponding pair-end reads were merged with PEAR v0.9.11 (Zhang et al., 2014) by specifying quality threshold to 26. After concatenating all sequences into a single FASTQ file, the sequences underwent a series of quality filtering with VSEARCH v2.14.1 (Rognes et al., 2016), including 1) quality filtering by limiting the maximum number of expected errors per read to 1; 2) frequency denoising with default alpha diversity parameter and minimum size threshold as 4 respectively; 3) length filtering to remove all reads shorter than 300bps for Uni18S assay and 100bps for remaining assays respectively, and 4) chimera filtering to remove any potential chimera generated in the reads. The filtered sequences were then clustered into operating taxonomic units (OTUs) with VSEARCH and a 97% similarity threshold greedy clustering method. The comprised OTU sequences were subsequently mapped to samples with VSEARCH (Rognes et al., 2016) to generate corresponding read table that showed the OTUs found in each sample and their corresponding read counts. Dietary metabarcoding is prone to co-amplification of secondary prey i.e. taxa presented in the diet of prey organism, that could create fabricate prey taxa and artificially increase prey richness (Deagle et al., 2019). Therefore, we applied a minimum sequence copy thresholds to each assay following the proportional method from Drake et al. (2022) to remove rarer sequences with relatively low read counts, and also low-level background noises from secondary predation and PCR/sequencing bias to improve data accuracy. Briefly, we remove any read counts less than a predefined proportion of the total read counts in each assay based on their respective sequencing depth. The proportion threshold 0.1% for Uni18S assay, 0.01% for both Fish16S and Ceph18S assay, and 0.001% for Crust16S assay respectively. We also removed any read counts within each OTU that were lower than the counts found in the negative controls.

The generated OTUs were searched against a local copy of GenBank nt database (downloaded on 19th November 2022) from National Center for Biotechnology Information (NCBI; Sayers et al., 2022) using BLASTn v2.12.0 with e-value of 0.001 (Altschul et al., 1990). The BLAST results were parsed through the python script taxonomy_assignment_BLAST_V1.py (Joseph7e, 2020) in order to assign final taxonomy based on the top BLAST hits and best consensus taxonomy that match the provided identity percent cut-offs. The minimum percent identity cut-offs for species and family were set to 98% and 91% respectively. OTUs that represented terrestrial fauna and flora, fungi, protozoans, dinoflagellates, algae, heterokont, diatom, zooplanktons, flatworms, and parasites were considered as non-prey items or environmental contaminants and removed from the dataset. Taxonomic nomenclature was based on the World Register of Marine (WoRMS Editorial Board, 2023)

### Statistical analyses

All statistical analyses and figure plotting were performed in R version 4.2.2. The influence of annealing temperature and blocking primer concentrations on rank transformed Cq values in each of the host species were analysed with Kruskal-Wallis rank-sum test and Dunn test. The potential effects of blocking primers on potential prey species amplifications were also analysed by Mann-Whitney U-Test.

Read tables were transformed into occurrence data to show the presence/ absence records of taxa across samples. As the taxa-specific sequence recovery bias of the primer assays used in this study had not been fully investigated (e.g. feeding trials), therefore read abundance data such as relative read abundance (RRA) that utilized read counts was not incorporated in this study. RRA also has issues in propagating the potential error in the dataset when incorporating reads from multiple primer assays contrary to occurrence data (Deagle et al., 2019). The blocking primer in 16S Fish assay also showed differential effects on amplification efficiency on different prey species (Figure S2), in which such disparity was also found to have significant effects on nontarget species relative read abundance(Piñol et al., 2015). Therefore, occurrence data is demonstrated to be the more appropriate metric to summarize sequence data in this multi primer assay study. After removing the low-level background noise through the minimum sequency copy threshold in previous step as suggested by Deagle et al. (2019) for occurrence data in dietary studies, the final taxa and read table were summarized as percentage of occurrence (POO) for each taxon in a sample, and frequency of occurrence (FOO) that showed the proportion of samples in which a taxon was detected for downstream analyses.

Individual sample completeness for each primer assay were indicated by rarefaction analysis with rarecurve function, and species accumulation curve analysis for each host species were done by specaccum function that were both found in vegan version 2.6-4 (Oksanen et al., 2008). Number of taxa found in each sample were used to imply taxa richness to evaluate and compare between different host species through ANOVA and Tukey HSD test. The overlapping prey taxa between the host species were visualized with Venn diagram by InteractiVenn virtual tool (Heberle et al., 2015). Heatmap of FOO of prey taxa across host species and associated hierarchical clustering were performed in R package Complexheatmap v 2.15.4 (Gu, 2022). The degree of prey taxa differentiation between host species were quantified with package Betapart v1.6 (Baselga & Orme, 2012), where the overall diversity was measured by Sørenson’s dissimilarity index (β_sor_), which was further partitioned into the Simpson dissimilarity index (β_sim_) to infer degree of prey taxa replacement between host species, and the nestedness resultant dissimilarity index (β_sne_) to infer prey taxa nestedness between host species.

The influence of host species, digestion level in stomach, fish total length, fish weight, and sampling season on recovered prey taxa richness were tested by performing PERMANOVAs with the function Adonis2 (vegan; Oksanen et al., 2008) coupled by Jaccard dissimilarity index and 999 permutations. The interaction between host species and any significant factor were then further investigated with Pairwise PERMANOVAs. Permutational analysis of multivariate dispersions (PERMDISP) was used to test homogeneity of variance within the host species with the function betadisper (vegan; Oksanen et al., 2008). The patterns of dietary taxa composition across host species were examined through nonmetric multidimensional scaling (nMDS) ordination based on Jaccard dissimilarity distance matrix, which were performed and visualised using metaMDS in vegan (Oksanen et al., 2008). Similarity percentage (SIMPER) was used to determine the contribution of individual prey taxa to beta diversity between the diet of species pairs. Function simper.pretty from R script simper_pretty.R was used to perform SIMPER on all species pairs, and the results from it were import into another function Kruskal.pretty from R script R_krusk.R to evaluate the respective statistical significance using Kruskal-Wallis rank-sum test with false discovery rate (FDJ) p-value corrections (Asteinberger9, 2020). The percentage cutoff and low cutoff value were set as 1 and 0.08 respectively.

## Results

### Sample Collection

A total of 63 fish were collected in the entire sampling period (Table S1). Overall, the proportion of samples with empty stomach with no prey remains i.e. digestion level 4 is 71.4%. The wet weight, total length (TL) and digestion level were found to significantly differ between species (Table S2; Kruskal-Wallis rank-sum test; *p* < 0.0001). Subsequent post hoc Dunn’s test with Bonferroni method to adjust p-value indicated that the differences were mostly caused by TGGG, which found to be significantly heavier and longer in terms of TL than *E. awoara*, *E. bleekeri* and *E. quoyanus* (p < 0.001). For digestion level, *E. quoyanus* had significantly lower levels than E. awoara (p < 0.001), *E. coioides* (p = 0.01) and TGGG (p < 0.001), while the digestion level of TGGG was also found to be significantly higher than that of *E. bleekeri* (p=0.01). Despite the overall low levels of identifiable prey (stomach content with digestion level 1 for visual identification) with only 6 out of 63 samples, the metabarcoding approach allowed disclosure of prey from 75% of all samples.

### Prey-specific high-throughput sequencing

Blocking of host species DNA was successful, with a minimum 10 cycle increase in Cq values regardless the annealing temperature or blocking primer to Fish16S primer ratio (Figure S1). Furthermore, the blocking primers showed minimal effect on the prey species (Figure S2; Mann-Whitney U-Test; *p*> 0.05) which shows that the designed blocking primers have high specificity towards targeted host species.

38,674,496 reads from 63 Fish16S assay samples, 10 Ceph18S assay samples, 29 Crust16S assay samples, 65 Uni18S assay samples and 2 PCR negative control samples remained after quality filtering and removal of non-prey taxa reads. Reads per sample range from 1,009 to 1,293,367 in Fish16S assay, 385 to 1,423,220 in Ceph18S assay, 323 to 2,008,225 in Crust16S assay and 116 to 53,517 in Uni18S assay respectively. The final number of reads per sample varied between individuals without systematic trends between host species (Kruskal-Wallis rank-sum Test, chi-square = 6.31, df = 4, p = 0.177). Six *E. awoara* samples and eleven TGGG samples had zero taxa or only host species detected, which all except one *E. awoara* had empty stomachs, and were removed from subsequent further analyses. The final number of samples having prey taxa detected was 36 (57.1% of all samples) in Fish16S assay, 7 (11.1%) in Cepha18S assay, 25 (39.7%) in Crust16S assay, and 10 (15.9%) in Uni18S assay respectively. The rarefaction curves of each assay showed a plateau was reached in all cases (Figure S3), indicated the sequencing depth was adequate to capture the number of OTUs presented in samples. On the other hand, the OTUs accumulation curve did not reach a plateau for all host species respectively (Figure S4), indicated more sampling is required in order to sample the full coverage of prey taxa.

For the final dataset after taxonomic assignment, 271 OTUs were considered as prey taxa which consisted of 61 unique prey taxa. 30 of which (49.2%) were assigned to species level, 11 (18.0%) to genus level, 17 (27.9%) to family level and 3 (4.91%) to order level. These taxa were consisted of 3 phylum namely Chordata, Arthropoda and Mollusca, which further categorized into 21 different orders. 32 (66.7%) intestine samples were found with prey OTUs despite absence of prey in stomach content. Nonetheless, there is a correlation between number of prey taxa and different digestion level (Kruskal-Wallis rank-sum test, Chi-Squared value = 19.3, p < 0.001). Subsequent post hoc Dunn’s test with Bonferroni adjusted p value showed samples with digestion level 4 had significant lower number of prey taxa than samples with digestion level 1 (p = 0.02) and level 2 (p = 0.003) respectively.

### Description and Comparison of diet

Number of prey taxa ranged from 1 to 9 found among the samples, with TGGG having the lowest count of prey taxa in general. There were significant differences in mean number of prey taxa between host species (Krustal-Wallis test, Chi-Squared value = 13.9, p=0.007). Subsequent post hoc Dunn’s test with Bonferroni adjusted p-value showed there is significant higher number of prey taxa found in *E. quoyanus* than in TGGG (p = 0.009; Figure 2a). No prey taxa were found to be shared across all 5 host species and only a few prey taxa are common across four of the host species, including two teleosts taxa namely Siganus and Lutjanidae, as well as two Crustacea taxa Portunidae and *Charybdis helleri*, and lastly one cephalopod Sepiolidae (Figure 2b). Only few prey taxa were at high frequency of occurence (FOO) among the host species. Prey taxa that had 50% or above FOO included *Omobranchus punctatus* in TGGG (68.8%), *Pleuronichthys Cornutus* in *E. coioides* (66.7%), Portunidae (62.5%) in *E. quoyanus*, *Charybdis hellerii* in *E. bleekeri* (66.7%) and *E. quoyanus* (62.5%), and lastly *Thalamita Admete* and decapoda in *E. quoyanus* (both 50.0%) (Figure 3).

**Figure 2.**
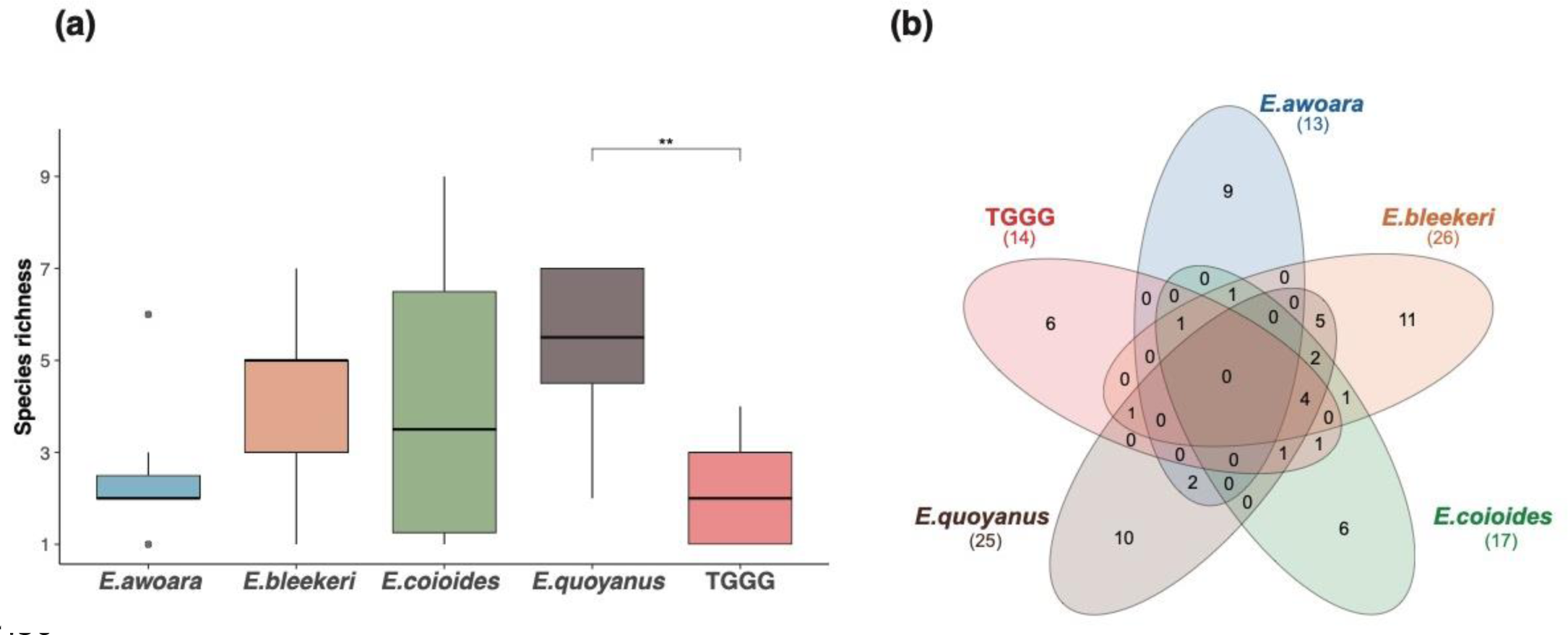
(a) Distribution of prey taxa number (taxa richness) found in each host species. Asterisks above represent significant pairwise mean differences between species indicated from post hoc Dunn’s test; (b) Number and overlap of prey taxa between the host species.

**Figure 3.**
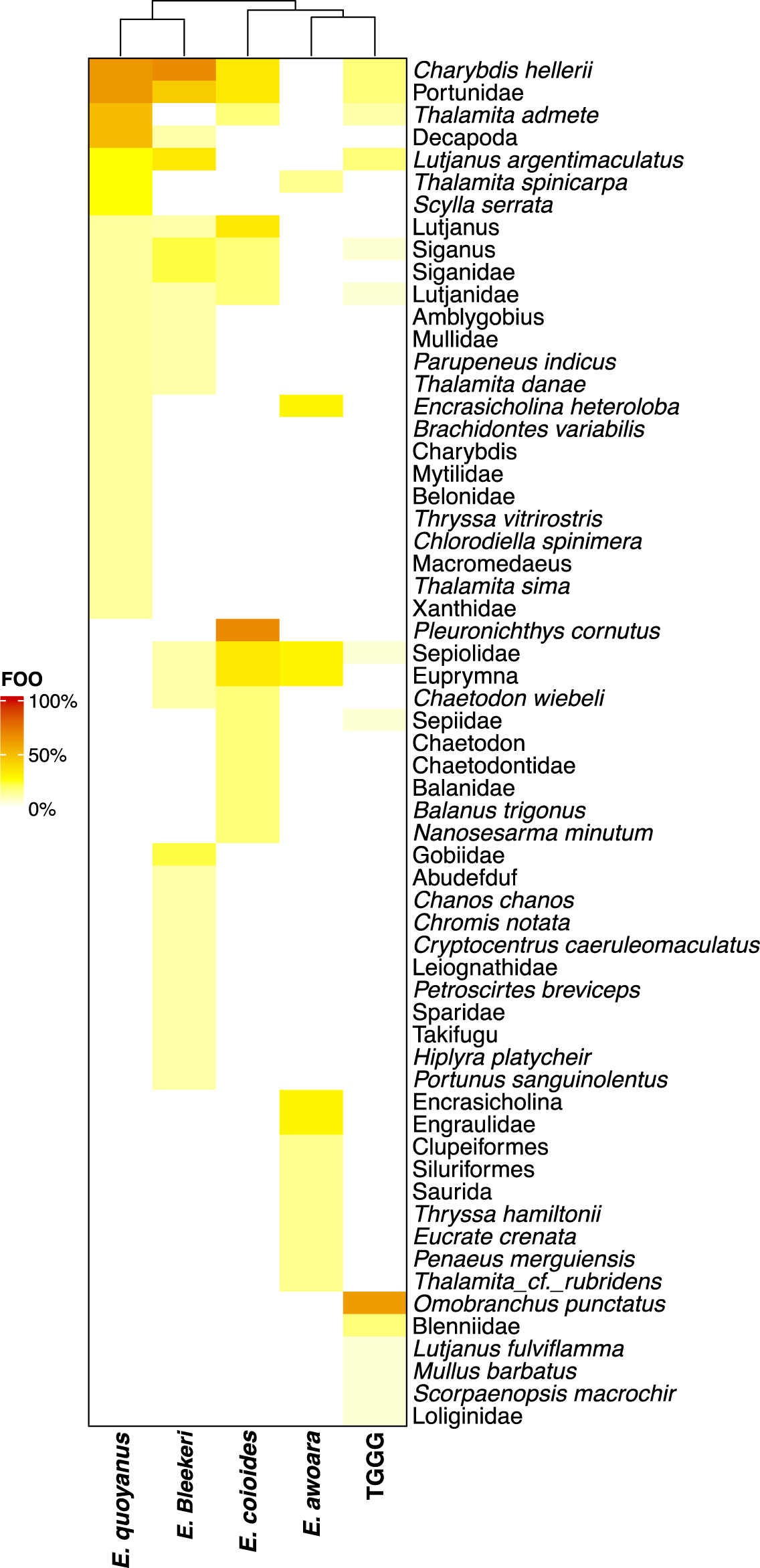
Frequency of Occurrence (FOO) of prey taxa found in samples of all 5 host species.

On the other hand, there are a number of unique prey taxa, with 42 only found in one of the extracted host species. Within these taxa there were 6 prey taxa uniquely found in TGGG samples, including 5 teleosts namely Blennidae, *Omobranchus punctatus, Lutjanus fulviflamma, Scorpaenopsis macrochir* and *Mullus barbatus*, and 1 cephalopod namely Loliginidae. TGGG and *E. coioides* both had the fewest unique prey taxa among the host species, with 9 unique prey taxa found *in E. awoara*, 11 in *E. bleekeri*, and 10 in *E. quoyanus* respectively (Figure 2b). In terms of prey diversity, *E. bleekeri* had the most diverse prey taxa among all host species, with 12 different orders of prey taxa found in the samples while for *E. awoara* with the lowest number of order of prey taxa, only 5 orders were found. TGGG (8), *E. coioides* (7) and *E. quoyanus* (8) shared similar number of taxa.

A clear pattern of dietary partitioning across species was found. The diet composition of *E. quoyanus, E. coioides* and *E. bleekeri* were clustered together and separated from that of TGGG and *E. awoara* as observed in hierarchical cluster analysis (Figure 3). Analysis of intestine contents in terms of percentage of occurrence (POO) also showed a large variation between TGGG and other host species (Figure 4). Notably, order Blenniformes with high POO (42.4%) found in TGGG was absent in other host species. The same was observed in *E. awoara* where Clupeiformes were found with high POO (44.5%) but not in most other host species. POO of prey taxa profiles for *E. quoyanus* was dominated by Decaopda, while diet of *E. coioides* and *E. bleekeri* are generally similar, with Decapoda being the most frequently taxa, followed by other taxa such as Lutjaniformes and Acanthuriformes. In terms of Sørensen dissimilarity between TGGG and other host species, difference in prey taxa between TGGG and *E. awoara* were found to be highest (0.93), then *E. bleekeri* (0.70), *E. quoyanus* (0.69) and lastly *E. coioides* (0.55). For other species pairs, Sørensen dissimilarity was found to be highest between *E. awoara* and *E. bleekeri* (0.90), and lowest between *E. bleekeri* and *E. quoyanus* (0.53) respectively. Simpson dissimilarity i.e. species turnover contributed the most to beta diversity between species, accounted for over 90% of Sørensen dissimilarity in most species pairs (Table S3). PERMANOVA also showed similar results where diet compositions were found to be significantly different between the species (df = 4, F = 2.73, p = 0.001). Subsequent pairwise comparison demonstrated that significant diet composition differences were found in most species pairs associated with TGGG or *E. awoara* (Table 2).

**Figure 4.**
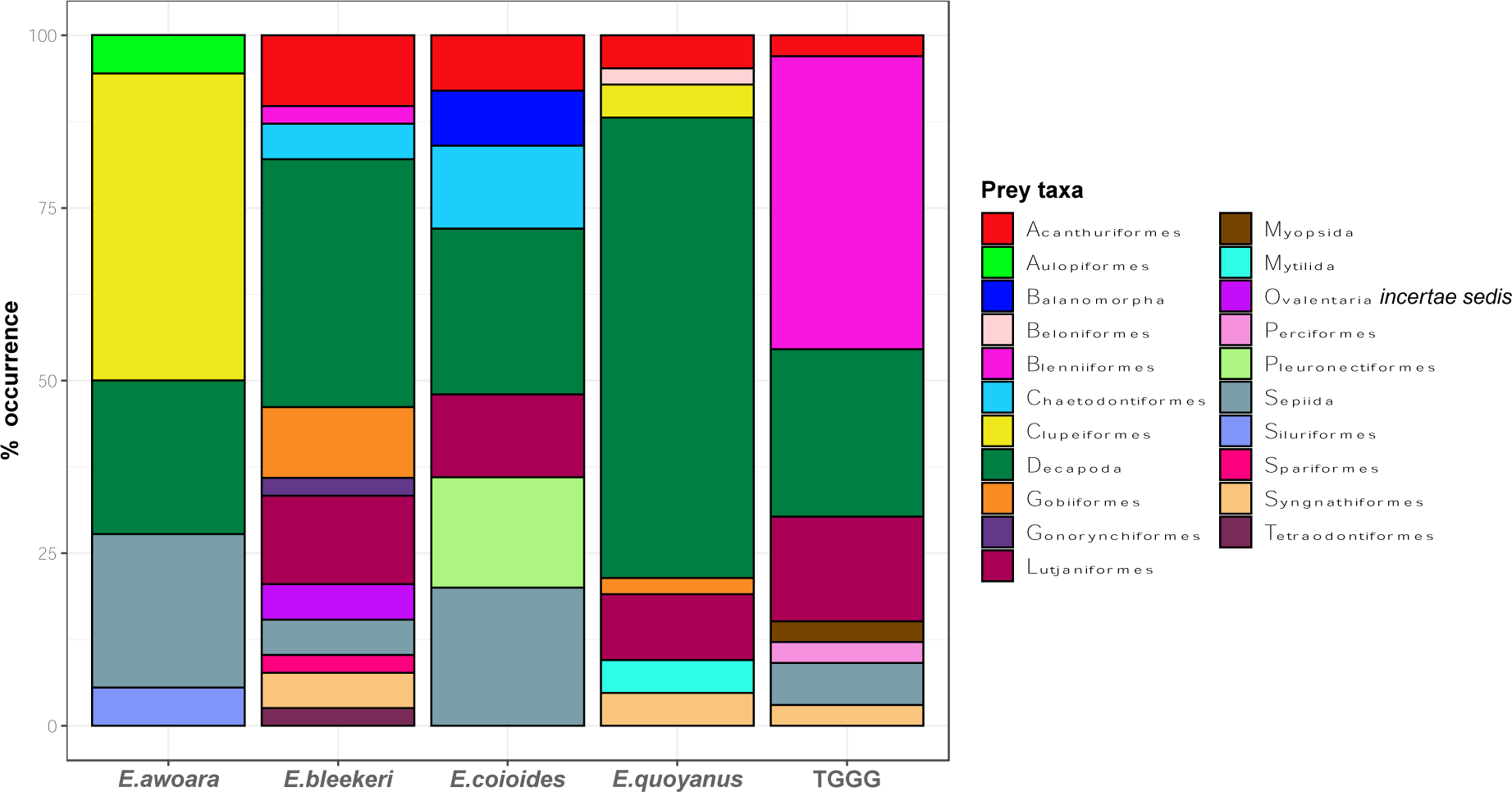
Percentage of Occurrence (POO) of prey taxa in each host species. Prey taxa were combined according to their associated Order.

**Table 2.** PERMANOVA pairwise comparison results of all species pairs. The final p-values were adjusted by Bonferroni correction.

| Species pair | F Model | Adjusted p value |
| --- | --- | --- |
| *E. bleekeri* / TGGG | 3.63 | 0.02 |
| *E. bleekeri* / *E. quoyanus* | 1.13 | 1 |
| *E. bleekeri* / *E. coioides* | 1.81 | 0.3 |
| *E. bleekeri* / *E. awoara* | 2.08 | 0.01 |
| TGGG / *E. quoyanus* | 4.12 | 0.02 |
| TGGG / *E. coioides* | 3.97 | 0.02 |
| TGGG / *E. awoara* | 3.88 | 0.01 |
| *E. quoyanus* / *E. coioides* | 2.29 | 0.03 |
| *E. quoyanus* / *E. awoara* | 2.57 | 0.01 |
| *E. coioides* / *E. awoara* | 1.72 | 0.67 |

While the diet differed among species, the interaction between species and season was also significant (PERMANOVA: df = 1, F = 1.42, p = 0.03). Significant inter-specific dietary partitioning was further identified between TGGG and E. awoara in dry season (Pairwise-PERMANOVA; p = 0.009), as well as between TGGG and *E. quoyanus* in wet season (Pairwise-PERMANOVA; p = 0.042) (Table 3). Other covariate factors including TL, weight and digestion level in stomach were not significant towards species diet composition as shown in PERMANOVA (p>0.05). All species showed homogenous dispersion in ordination space based on their centroids (PERMDISP: df = 4, F = 0.838, p = 0.509). NMDS analysis also revealed similar patterns in diet dissimilarity, where ellipses of TGGG and *E. awoara* did not overlap with others, while ellipses of *E. quoyanus, E. coioides* and *E. bleekeri* overlapped and clustered together (Figure 5; stress = 0.155). The ANOSIM result (R= 0.350, p = 0.001) further supported the significant dissimilarity in diet composition between species. Finally, SIMPER analysis together with Kruskal-Wallis rank-sum test identified a total of 6 prey taxa being most significant in discriminating between TGGG and the associated four distinct host species pairwise comparison. *Omobranchus punctatus* in TGGG contributed the most in all four pairwise comparison (11.6% to 17.6%; Table S4). *Charybdis helleri* in *E. bleekeri* (10.6%) and *E. quoyanus* (9.12%), Pleuronichthys cornutus in *E. coioides* (17.1%), Decapoda and Portunidae in *E. quoyanus* (8.80% and 9.46% respectively) also contributed significantly to their pairwise dissimilarities associated with TGGG.

**Figure 5.**
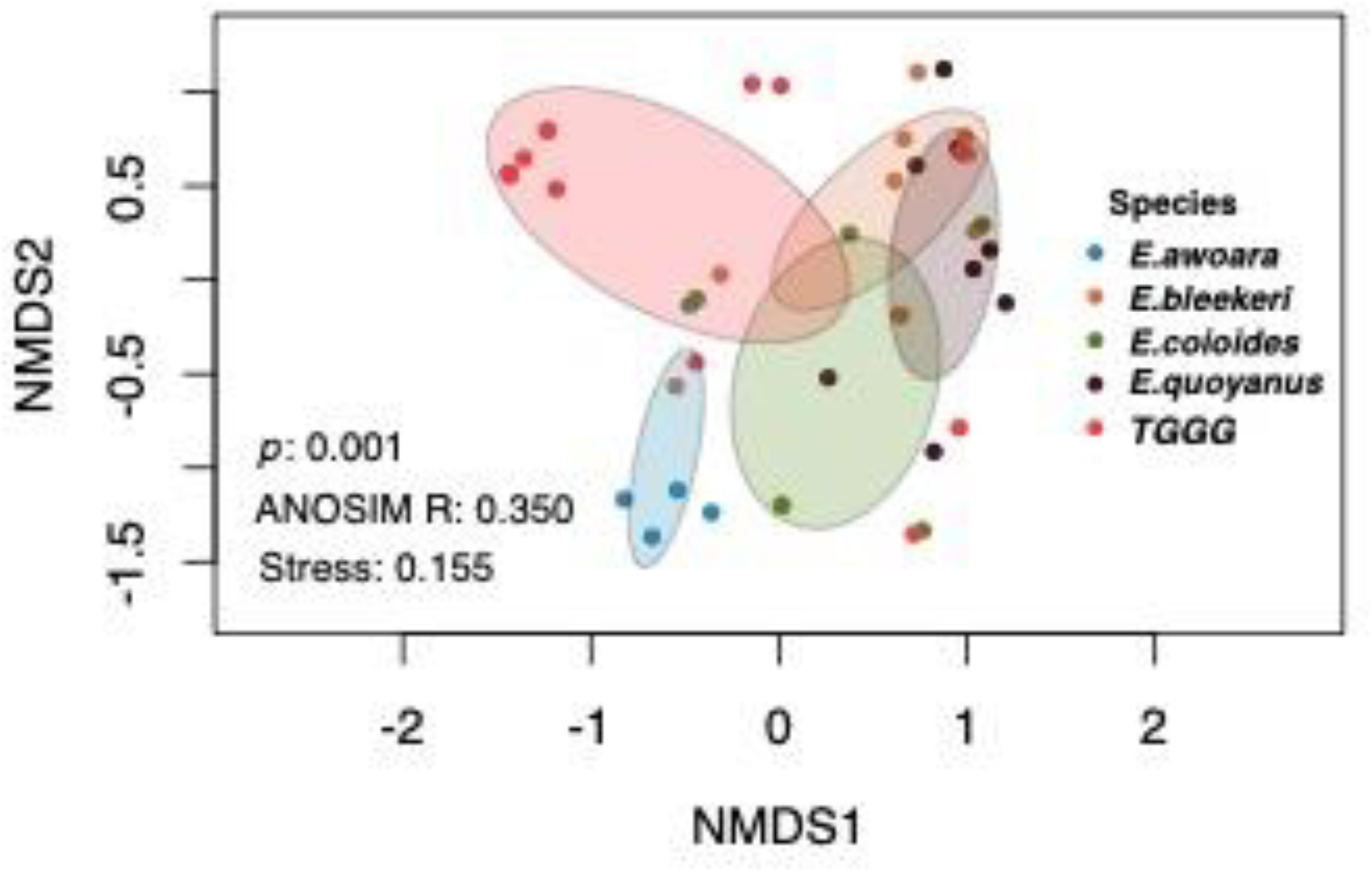
Multidimensional scaling analysis of prey taxa detected in five host species (dots) based on Jaccard dissimilarities. Standard deviation with default confidence interval ellipses are shown per species.

**Table 3.** PERMANOVA pairwise comparison results of species pairs in each season. The final p value was adjusted by Bonferroni correction.

|  |  |  |
| --- | --- | --- |
| 2020 Dry Season |  |  |
| Species pair | F Model | Adjusted p value |
| TGGG / *E. coioides* | 2.54 | 0.063 |
| TGGG / *E. awoara* | 2.93 | 0.009 |
| *E. coioides* / *E. awoara* | 1.09 | 1 |
| 2021 Wet Season |  |  |
| Species pair | F Model | Adjusted p value |
| *E. bleekeri* / TGGG | 2.44 | 0.078 |
| *E. bleekeri* / *E. quoyanus* | 1.13 | 1 |
| *E. bleekeri* / *E. coioides* | 0.98 | 1 |
| TGGG / *E. quoyanus* | 2.82 | 0.042 |
| TGGG / *E. coioides* | 2.06 | 0.336 |
| *E. quoyanus* / *E. coioides* | 1.41 | 0.612 |

## Discussion

Dietary information and partitioning between native species and an introduced potential competitor can reveal the extent of overlapping resource use which is vital in understanding how anthropogenic pressures impact local trophic interactions and ecosystem dynamics. We used a contemporary molecular approach with fine-scale taxonomic resolution to unveil the diet of the artificial hybrid TGGG which is released into the wild by religious practise and a potential threat to the native fauna. We demonstrate the capability of TGGG to prey on a variety of prey taxa across teleost, crustaceans and cephalopods. Although groupers are generally perceived as opportunistic and versatile predators (Gibran, 2007), we found numerous unique prey taxa in TGGG and four other studied species (*Epinephelus* awoara, *E. bleekeri*, *E. coioides* and *E. quoyanus*), which displays patterns of dietary partitioning with clear differential prey preferences in a coastal ecosystem.

We reveal that the diet composition of TGGG is dissimilar from that of other *Epinephelus* spp., indicating that TGGG is utilizing different resources and occupying an unique dietary niche in the community. This finding do not coincide with our hypothesis, where we speculated higher overlap of dietary niche between TGGG and *Epinephelus* spp. as suggested from other farm escape examples (Nabaes Jodar et al., 2020). One possible explanation for such differences is that most dietary studies on farm escapes or *Epinephelus* spp. are based on visual inspection only and fail to give precise prey taxa identification compared to the metabarcoding approach in this study. The fact that we found diet partitioning among the hybrid and the native groupers may have several explanations. One possible reason may derive from the usage of different microhabitats, due to surface feeding on pellets inside aquaculture facilities (Nabaes Jodar et al., 2020), TGGG may proceed to conduct similar feeding strategies and therefore utilize different resources than other *Epinephelus* spp. in the wild. This is supported by a number of unique prey taxa in TGGG including a family of squid, Loliginidae, and *Omobranchus punctatus* which are frequently found in shallow and surface waters. Another explanation is related to the parental species of TGGG and inheritance of their associated ecological niche. As hybrids were found to have high dietary overlap with the parental species (Montanari et al., 2014), hybrids may utilize similar resources in the ecosystem and share dietary niche to a certain degree. TGGG may have inherited the unique niche from its parent and displayed dietary partitioning with other *Epinephelus* spp., however no dietary information exist for the two parental species. To pinpoint the exact reason why the diet of the hybrid is unique, further elaborate field observations would be necessary, including spatial and temporal resources utilization, as well as feeding behaviour of TGGG in the wild.

On the other hand, the diet of local *Epinephelus* spp. showed a high diversity of prey, with teleosts and crustaceans especially crabs being the major prey taxa, which aligned with previous visual analysis on diet composition of *E. coioides* and *E. quoyanus* in tropical reef or mangrove ecosystem (Connell, 1998; Kulbicki et al., 2005). Each species of *Epinephelus* however displayed distinct patterns of prey preferences, including differences in prey richness, frequencies and diversity. Morphological features of predators could be one main factor explaining these partitioning patterns observed (Wainwright & Richard, 1995). For groupers, body size and mouth opening size in terms of relative mouth width and height to the body length in *E. coioides* are found significantly altering prey diversity and stomach fullness (Chuaykaur et al., 2020), as larger body size and mouth opening provide greater suction force (Wainwright & Richard, 1995), enabling teleost predators to prey on wider range of prey. Yet our results showed no apparent correlation of body size and diet composition, implying there are also other factors impacting the differences in diet choices other than ecomorphological features. Other potential factors include utilization of different feeding behaviour or tactics that could introduce niche partitioning which could lead to subtle variation of resources consumed (Kent & Sherry, 2020) and enable co-existence of closely related species. A number of foraging behaviours have been recorded among the Serranid family to which *Epinephelus* species belong, such as ambush predation, patrolling, drift feeding (Gibran, 2007), cleaning behaviour (Hackradt et al., 2013) or even use body camouflage offensively to approach potential preys (Watson et al., 2014). Hence, it may be different behaviours or occupying different microhabitats that allow utilization a diverse range of preys among Serranidae species.

Compared with the diet composition of the local native *Epinephelus* spp., the TGGG hybrid fed on a significantly lower diversity of prey, along with the highest portion of samples with no prey taxa detected, as well as lowest median number of prey taxa across all host species. This suggests that TGGG released from mercy release practices, similar to cases of aquaculture farm escapes, showed variations between individuals in acquiring feeding behaviour to prey in the wild (Skilbrei et al., 2015). This indicated that a large portion of individuals in release events may likely die shortly upon release, while a few of them are capable of slowly acquiring feeding behaviour and feed in the wild efficiently to perform diet transitioning from farm food to wild prey. As frequent mercy release events keep introducing large numbers of TGGG into wild, there is therefore a potential for a thriving wild population despite low initial survival rate. Introduced marine teleosts can drive devastating changes in population dynamics and structure in the community, such as the invasion of Indo-Pacific Lionfish (Pterois spp.) into western Atlantic that led to the reduction in prey taxa richness, location extinctions and changes in local predator populations (Albins, 2013; Albins & Hixon, 2008; Ingeman, 2016; Lesser & Slattery, 2011). Cases of introduced grouper, such as *Cephalopholis argus*, a species from Indo-Pacific which was intentionally released in Hawaii in 1950s was successful at establishing populations and became a dominant piscivore in the reefs (Dierking et al., 2009). Furthermore, being an apex predator itself, along with the absence of other predators such as sharks that are long been locally extinct in the region (Cardeñosa et al., 2022), implies that there is essentially no restraint to TGGG population growth through top-down control in the food chain (Baum & Worm, 2009). The TGGG occupation on vacant dietary niches among local species suggests an additional potential of introduction success, as unoccupied niches are found to be linked with successful establishment of introduced species in aquatic ecosystem (González-del-Pliego et al., 2023; Tapkir et al., 2023). Together with evidence from laboratory settings that TGGG can produce viable F2 offspring (Ching et al., 2018), there is potential for the establishment of TGGG locally. Still, many factors will indeed affect the success of an invasive species such as fish sexual maturity and seasons of escape to play a role in the long-term survival (Nabaes Jodar et al., 2020; Skilbrei et al., 2015). To fully understand the long-time survival ability of the TGGG time spent since release into the wild throughout the seasons as well as other unknown ontogenetic dietary shifts would be necessary. Nonetheless, we show the possible capabilities of establishing wild populations in TGGG and hence the potential ecological impact brought by TGGG should not be underestimated.

It is noteworthy that despite TGGG being the most popular type of grouper hybrid, there are increasing number of other aquaculture hybrids available in the market which also inherent the massive body size from parents such as *E. lanceolatus* x *E. polyphekadion* and *E. moara* x *E. lanceolatus* (Chen et al., 2018; Rimmer & Glamuzina, 2019). These types of hybrids, although received far less attention, are also targets of mercy release and hold similar potentials as TGGG to pose threats to local ecosystems.

Our study presents critical information on wild released hybrids and show distinct resource utilization of introduced artificial hybrid TGGG and four other congeneric grouper species in a coastal species-rich ecosystem. Our results provide the first evidence that an artificial hybrid aquaculture fish could prevail in a coastal marine ecosystem. Rarely documented religious tradition are the main cause for the release of the hybrids, adding evidence for an understudied anthropogenic activities impacting and altering the trophic interactions in coastal marine ecosystem.

## Acknowledgements

We thank Chris Cheung, Jason, Tony, Mr. Leung and Mr. Ma on assisting sampling collection. Thank you Dr Allen To, Eric Keung, Ryan Cheng, Stan Shea and Yiu Wai Hong from 114°E Hong Kong Reef Fish Survey for providing underwater photographs to this manuscript. This work was supported by the Environmental and Conservation Fund (ECF 2020-142).

## Data availability

Raw sequence data files are available via dryad platform (https://datadryad.org/stash/share/zaMXw4_s6hYigjk3fBgJD2ZMuWUdbvxr_3icuVnbCrI).

## Notes

### Competing Interest Statement

The authors have declared no competing interest.

