## Supplementary data for "Distinct resource utilization by introduced man-made grouper hybrid: an overlooked anthropogenic impact from a longstanding religious practise"

### **Dietary composition and niche partitioning of introduced hybrid *Epinephelus fuscoguttatus* x *E. lanceolatus* and native grouper species**

Arthur Chung[1], Celia Schunter[1*]

[1]The Swire Institute of Marine Science and School of Biological Sciences, The University of Hong Kong, Pokfulam Road, Hong Kong

*

Supplementary Data

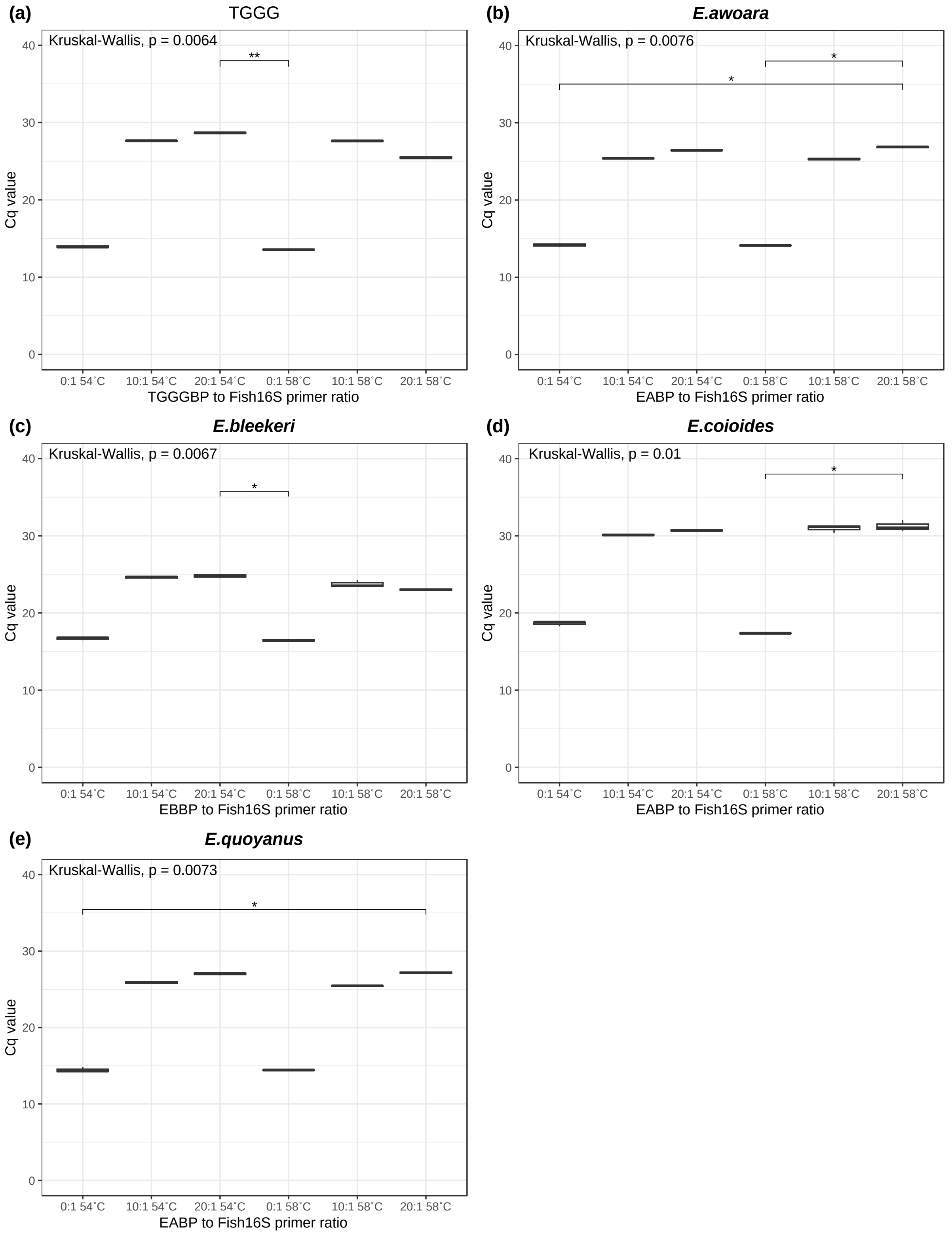

Figure S1. Mean Cq value of treatment on genomic DNA from (a) TGGG, (b) *E. awaora*, (c) *E. bleekeri*, (d) *E. coioides* and (e) *E. quoyanus* at different level of species-specific blocking primer to Fish16S assay primer ratio (0 to 1, 10 to 1 and 20 to 1) and different annealing temperature (54˚C and 58˚C). Asterisk above each bar represented significant pairwise mean differences between samples indicated from Dunn test.

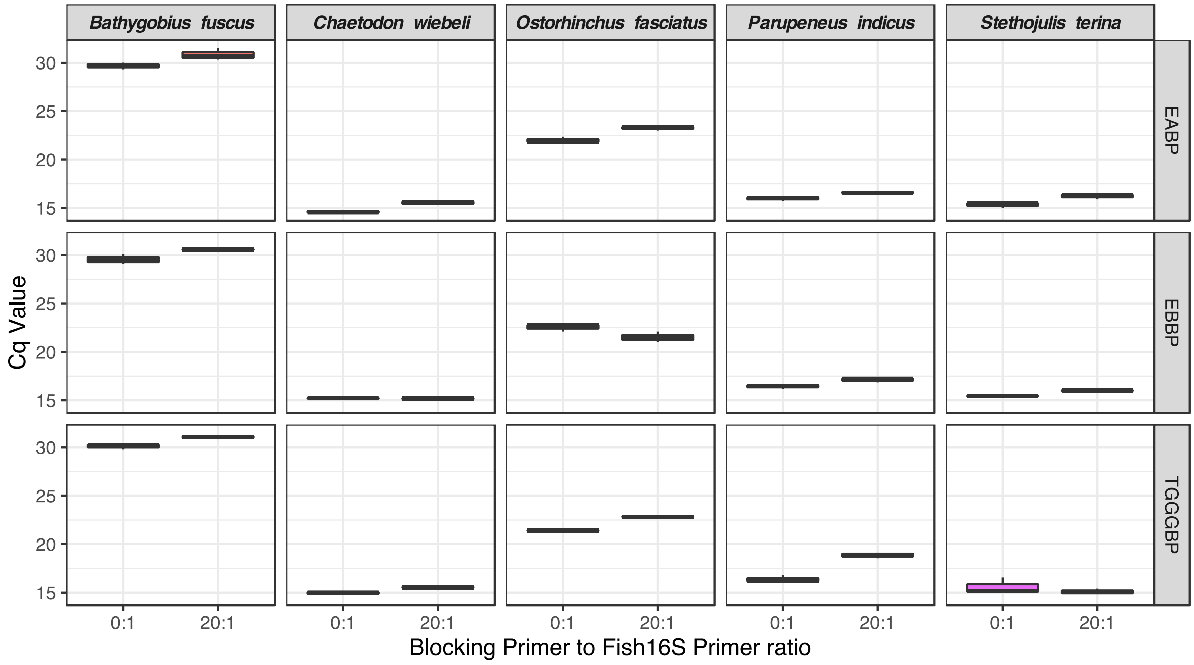

Figure S2. Mean Cq value of treatment on genomic DNA of different potential prey species, including *Bathygobius fuscus, Chaetodon wiebeli, Ostorinchus fasciatus, Parupeneus indicus* and *Stethojulis terina* at different level of species-specific blocking primer to Fish16S assay primer ratio (0 to 1 and 20 to 1).

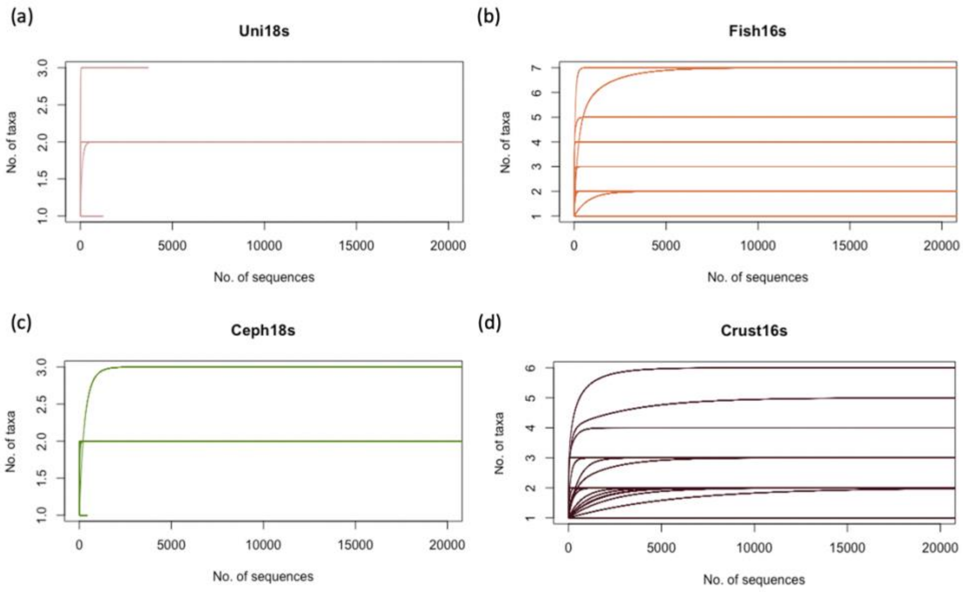

Figure S3. Rarefaction curve of samples from (a) Uni18S assay, (b) Fish16S assay, (c) Ceph18S assay and (d) Crust16S assay.

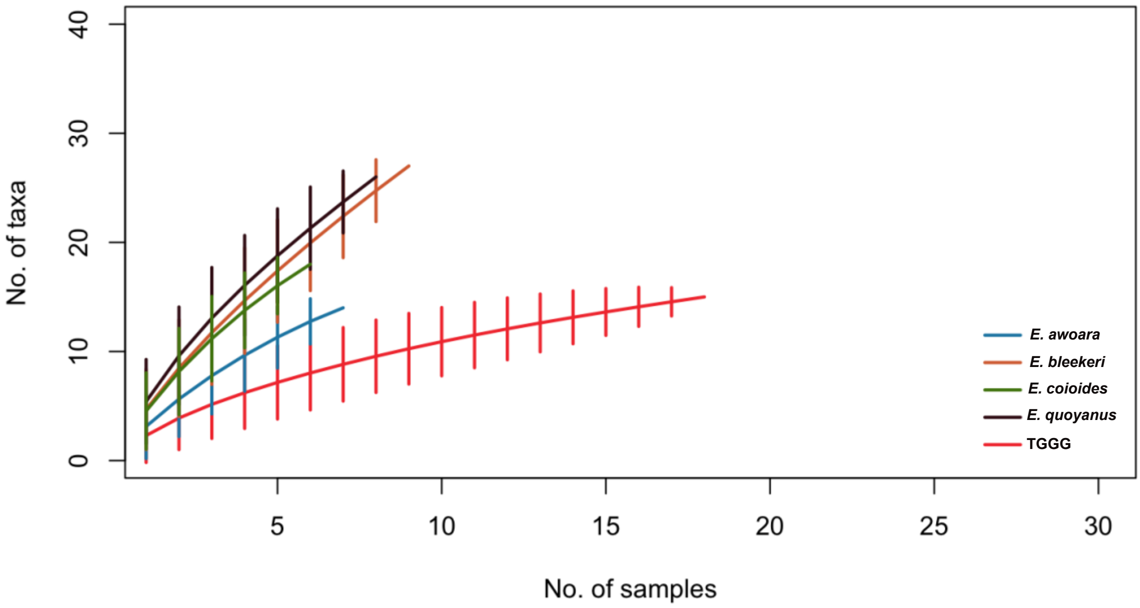

Figure S4. Accumulation curve showing the cumulative number of taxa being found in each of the host species as a function of number of samples. Error bars represented standard deviation.

Table S1. The TL, wet weight and digestion level of the collected samples from all five host species

| Species | Sample size (n) | TL (cm) (± SD) | TL min max | Wet Weight (g) (± SD) | Weight min max | Digestion level (Percentage) |
|---|---|---|---|---|---|---|
| TGGG | 27 | 34.1±9.27 | 26.4, 68 | 1108±1527 | 384, 6480 | 1: / (0%)<br>2: 1 (3.7%)<br>3: 2 (7.4%)<br>4: 24 (88.9%)) |
| *E. awoara* | 13 | 25.1±1.87 | 21.6, 27.1 | 282±61.2 | 172, 369 | 1: / (0%)<br>2: / (0%)<br>3: 2 (15.4%)<br>4: 11 (84.6%) |
| *E. bleekeri* | 9 | 22.8±3.60 | 18.9, 28.5 | 205±110 | 102, 385 | 1: 1 (11.1%)<br>2: 5 (55.6%)<br>3: / (0%)<br>4: 3 (33.3%) |
| *E. coioides* | 6 | 28.6±10.4 | 19.3, 44.9 | 469±527 | 109, 1483 | 1: / (0%)<br>2: 1 (16.7%)<br>3: / (0%)<br>4: 5 (83.3%) |
| *E. quoyanus* | 8 | 24.2±1.05 | 22.8, 25.6 | 243±40.4 | 173, 303 | 1: 5 (62.5%)<br>2: 2 (25%)<br>3: / (0%)<br>4: 1 (12.5%) |

Table S2. Results of Kruskal-Wallis test to examine TL, wet weight and digestion level difference between the host species

| Variable | df | Chi-squared value | p value |
|---|---|---|---|
| TL | 4 | 36.5 | <0.001 |
| Wet weight | 4 | 40.7 | <0.001 |
| Digestion level | 4 | 30.8 | <0.001 |

Table S3. Beta diversity between species pairs as calculated by betapart package in R

| | | | | |
|---|---|---|---|---|
| Simpson dissimilarity | | | | |
| | TGGG | *E. awoara* | *E. bleekeri* | *E. coioides* |
| *E. awoara* | 0.92 | | | |
| *E. bleekeri* | 0.57 | 0.85 | | |
| *E. coioides* | 0.50 | 0.85 | 0.47 | |
| *E. quoyanus* | 0.57 | 0.85 | 0.52 | 0.59 |
| Nestedness-resultant dissimilarity | | | | |
| | TGGG | *E. awoara* | *E. bleekeri* | *E. coioides* |
| *E. awoara* | 0.00 | | | |
| *E. bleekeri* | 0.13 | 0.05 | | |
| *E. coioides* | 0.05 | 0.02 | 0.11 | |
| *E. quoyanus* | 0.12 | 0.05 | 0.01 | 0.08 |
| Sørensen dissimilarity | | | | |
| | TGGG | *E. awoara* | *E. bleekeri* | *E. coioides* |
| *E. awoara* | 0.93 | | | |
| *E. bleekeri* | 0.70 | 0.90 | | |
| *E. coioides* | 0.55 | 0.87 | 0.58 | |
| *E. quoyanus* | 0.69 | 0.89 | 0.53 | 0.67 |

Table S4. Similarity Percentage (SIMPER) analysis on the contribution of individual taxa (Significance determined by Krustal-Wallis rank-sum test with false discovery rate (FDJ) p-value corrections) on diet partitioning between pairwise comparison of host species

| | | **SIMPER results (%)** | | | | | |
|---|---|---|---|---|---|---|---|
| **Species 1** | **Species 2** | **Decapoda** | ***Pleuronichthys cornutus*** | **Encrasicholina** | ***Omobranchus punctatus*** | **Portunidae** | ***Charybdis hellerii*** |
| **TGGG** | ***E. awoara*** | | | 8.99 | 17.6 | | |
| **TGGG** | ***E. bleekeri*** | | | | 14.0 | | 10.6 |
| **TGGG** | ***E. coioides*** | | 17.1 | | 16.4 | | |
| **TGGG** | ***E. quoyanus*** | 8.80 | | | 11.6 | 9.46 | 9.12 |
| ***E. bleekeri*** | ***E. awoara*** | | | | | | 9.67 |
| ***E. bleekeri*** | ***E. coioides*** | | 12.0 | | | | |
| ***E. coioides*** | ***E. awoara*** | | 15.6 | | | | |
| ***E. coioides*** | ***E. quoyanus*** | | 9.99 | | | | |
| ***E. awoara*** | ***E. quoyanus*** | | | | | 8.72 | 8.34 |
